# Comparison of three different eye-tracking tasks for distinguishing autistic from typically developing children and autistic symptom severity

**DOI:** 10.1101/547505

**Authors:** Juan Kou, Jiao Le, Meina Fu, Chunmei Lan, Zhuo Chen, Qin Li, Weihua Zhao, Lei Xu, Benjamin Becker, Keith M Kendrick

## Abstract

Altered patterns of visual social attention preference detected using eye-tracking and a variety of different paradigms are increasingly proposed as sensitive biomarkers for autism spectrum disorder. However, few eye tracking studies have compared the relative efficacy of different paradigms to discriminate between autistic compared with typically developing children and their sensitivity to specific symptoms. To target this issue, the current study used three common eye tracking protocols contrasting social versus non-social stimuli in young (2-7 years old) Chinese autistic (n = 35) and typically developing (n = 34) children matched for age and gender. Protocols included dancing people *vs*. dynamic geometrical images, biological motion (dynamic light point walking human or cat) *vs*. non-biological motion (scrambled controls) and child playing with toy *vs*. toy alone. Although all three paradigms differentiated autistic and typically developing children, the dancing people versus dynamic geometry pattern paradigm was the most effective, with autistic children showing marked reductions in visual preference for dancing people and correspondingly increased one for geometric patterns. Furthermore, this altered visual preference in autistic children was correlated with the ADOS social affect score and had the highest discrimination accuracy. Our results therefore indicate that decreased visual preference for dynamic social stimuli may be the most effective visual attention-based paradigm for use as a biomarker for autism in Chinese children. Clinical trial ID: NCT03286621 (clinicaltrials.gov); Clinical trial name: Development of Eye-tracking Based Markers for Autism in Young Children.

**Lay summary:** Eye-tracking measures may be useful in aiding diagnosis and treatment of autism, although it is unclear which specific tasks are optimal. Here we compare the ability of three different social eye-gaze tasks to discriminate between autistic and typically developing young Chinese children and their sensitivity to specific autistic symptoms. Our results show that a dynamic task comparing visual preference for social (individuals dancing) versus geometric patterns is the most effective both for diagnosing autism and sensitivity to its social affect symptoms.

## Introduction

Autism Spectrum Disorder (ASD) comprises symptoms including social communication deficits and unusual repetitive and restrictive sensory-motor behaviors displayed from an early age [Kanner, 1943]. The prevalence of ASD has increased to 1/59 according to the Centers for Disease Control (CDC) in the United States [Christensen et al., 2016], although estimates vary considerably worldwide with 1/38 in South Korea [Kim et al., 2011] and 1/255 in China [Wang et al., 2018]. While ASD twin studies have demonstrated a strong genetic contribution to ASD with heritability estimates ranging between 64% and 91% [Tick, Bolton, Happé, Rutter, & Rijsdijk, 2016], no reliable genotype biomarker is current available. Thus, the current ASD diagnostics rely entirely on the assessment of behavioral symptoms based on two domains (social communication and restricted and repetitive or unusual sensory behaviors) [Lord, Elsabbagh, Baird, & Veenstra-Vanderweele, 2018]. Among all the tools used for diagnosing of ASD is the Autism Diagnostic Observation Schedule [ADOS-2; Lord, Luyster, Gotham, & Guthrie, 2012; Lord et al., 2012] which is regarded as the “gold standard” measure of observational assessment [Kanne, Randolph, & Farmer, 2008] and used in both clinical and research fields. The symptoms of social interest deficits (eg. social eye contact, social facial expressions and social motion) have been treated as important hallmarks of autism and used as diagnostic criteria in ADOS-2 [Lord et al. 2012] as well as questionnaire-based assessments such as the Autism Diagnostic Interview [ADI-R-Rutter, Le Couteur, & Lord, 2003] and the social responsiveness scale [SRS - Constantino & Gruber 2012]. However, simple, sensitive, objective and quickly performed physiological measures of social interest deficits are required to increase diagnostic accuracy and evaluate the efficacy of novel treatment approaches.

Given the symptoms of social interest deficits in ASD, eye-tracking technology in particular has facilitated research in investigating altered social attentional biases in autistic individuals of all ages. One of the first studies to use this approach demonstrated that adolescent participants with ASD watching movies spent significantly less fixation time on the people shown and more on the background scene [Donald, 2002]. Subsequent studies have further developed the use of eye-tracking and different paradigms to establish robust objective and quantitative biomarkers for ASD compared with typically developing (TD) controls primarily by contrasting visual attention towards social as opposed to non-social stimuli. Overall, they have shown that individuals with ASD tend to show reduced attentional preference for social stimuli [Falck-Ytter, Bölte, & Gredebäck, 2013; Fujisawa, Tanaka, Saito, Kosaka, & Tomoda, 2014; Klin, Lin, Gorrindo, Ramsay, & Jones, 2009; Moore et al., 2018; Pierce, Marinero, Hazin, Mckenna, & Barnes, 2015; Shi et al., 2015] and biological motion [Atkinson, 2009; Jones et al., 2011; Koldewyn, Whitney, & Rivera, 2010; Murphy, Brady, Fitzgerald, & Troje, 2009; Rutherford & Troje, 2012; Saygin, Cook, & Blakemore, 2010]. However, not all studies have reported positive results, particularly those involving attention to biological motion, and conclusions are often hampered by small sample sizes [Atkinson, 2009; Jones et al., 2011; Koldewyn, Whitney, & Rivera, 2010; Murphy, Brady, Fitzgerald, & Troje, 2009; Saygin, Cook, & Blakemore, 2010].

Few eye-tracking studies have compared the relative efficacy of different paradigms contrasting preference for social and non-social stimuli and the majority have also primarily involved Caucasian children. In the current study we have therefore compared the efficacy of three different protocols for firstly discriminating between young Chinese children with ASD compared with TD controls and secondly their sensitivity to ASD symptoms quantified by ADOS-2. The three protocols chosen were firstly attention to dynamic geometric patterns versus individual dancing humans [Pierce et al., 2015; adapted to incorporate Chinese stimuli]; secondly a classical biological motion protocol (dynamic light point walking human (or cat) versus a scrambled control version) [Rutherford & Troje, 2012] and thirdly an adaptation of protocols [Chevallier et al., 2015; Frazier et al., 2018] investigating attentional bias towards children playing with a toy as opposed to the toy itself.

## Methods

### Ethics Statement

The experiment was approved by the Institutional Review Board, University of Electronic Science and Technology of China. The caregivers of participants provided written informed consent before study enrollment.

### Participants

77 young children were enrolled in the study. 43 of the children were diagnosed as having autism spectrum disorder (ASD) by qualified clinicians and without co-morbid psychiatric or neurological disorders were recruited from a local (Southwest Children’s Hospital in Chengdu) children’s hospital. The clinical diagnosis of ASD was based on either DSM-IV [APA, 1994] or ICD-10 [WHO, 1994]. However, additionally we used the Autism Diagnostic Observation Schedule, second edition (ADOS-2, Module T, 1, or 2) [Lord et al. 2012] to assess whether all ASD children met the criteria for autism and 8 children did not and were therefore excluded resulting in a total of 35 children in the ASD group. The ADOS-2 assessment was performed by two certificated research reliable researchers (J.K., J.L.). The 34 TD children were recruited from a local kindergarten by advertisement. The age and gender of the final ASD and TD groups were matched. All children, including the TD group, were assessed using a series of questionnaire-based measures including the Social Responsiveness Scale 2nd edition (SRS-2) [Constantino & Gruber 2012], Repetitive Behavior Scale Revised-2 (RBS-2) [Lam & Aman, 2007], Caregiver Strains Questionnaire [Brannan et al., 1997] (CSQ), Social Communication Questionnaire [Rutter et al., 2003] (SCQ) completed by one of their caregivers. Table 1 shows demographic and questionnaire scores for the ASD and TD groups and ADOS-2 scores for the ASD group. The study was pre-registered as a clinical trial at Clinical Trials.gov (NCT03286621).

**Table 1.** Demographics, questionnaire and ADOS scores for ASD and TD groups

| Characteristic | ASD<br>n=32 | TD<br>n=34 | *p* | Cohen's<br>*d* |
| --- | --- | --- | --- | --- |
| Age(year) | 3.72±1.25 | 4.10±0.47 | 0.114 | 0.41 |
| Male/Female | 26/6 | 25/9 | 0.454* | - |
| SRS-2 total score | 85.81±14.57 | 40.09±14.38 | <0.001 | 3.21 |
| RBS-2 | 22.22±13.86 | 13.15±14.30 | 0.012 | 0.66 |
| CSQ | 55.10±13.37 | 28.33±6.58 | <0.001 | 2.60 |
| SCQ | 17.47±5.28 | 22.85±4.65 | <0.001 | 1.10 |
| ADOS-2 Total Score | 18.94±4.10 | - |  |  |
| SA | 14.41±3.10 | - |  |  |
| RRB | 4.53±2.27 | - |  |  |

SRS = social responsivity scale; CSQ = Caregiver Strains Questionnaire; SCQ = Social Communication Questionnaire; RBS = Repetitive behavior scale; ADOS = Autism diagnosis observation schedule; SA = Social Affect component of ADOS and RRB Restricted and Repetitive Behavior component of ADOS. \* indicates Chi Square Test Value $\chi^2 = 0.560$ , $p = .454$

### Task Paradigms

Three eye-tracking paradigms were included in this study. The first paradigm is a *Dynamic Visual Preference Task (Task 1)* displaying dynamic dancing Chinese human versus dynamic geometry patterns. The second paradigm is an *Abstract Animate Point Visual Exploration Task (Task 2)* displaying point-light animate (walking human or cat) and inanimate (randomly moving point-light) videos. The last paradigm is a *Static Visual Preference Task (Task 3)*, comparing attention to static pictures showing a toy alone or with a child playing with it (Figure 1).

**Fig.1.**
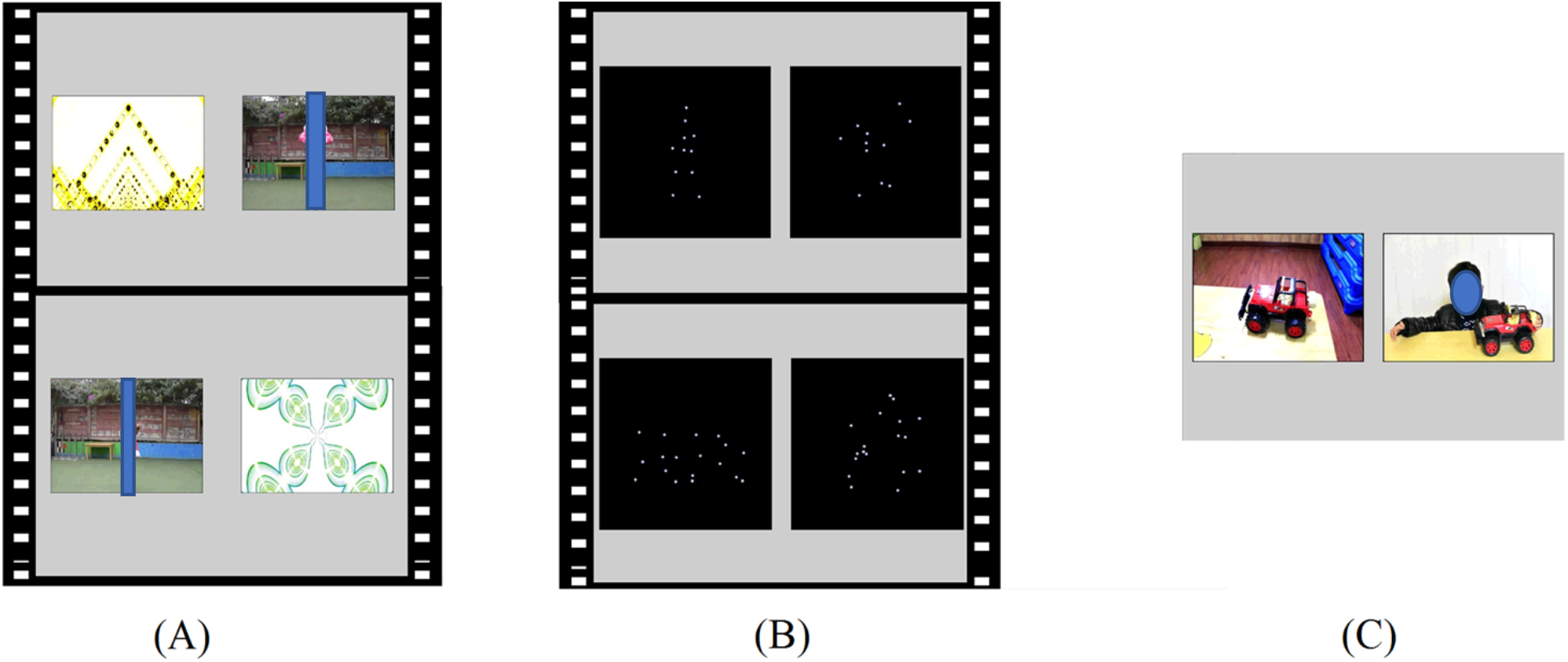
Examples of stimuli for each of the tasks: (A) Task 1: Dynamic Visual Preference Task (B) Task 2: Abstract Animate Point Visual Exploration Task (C) Task 3: Static Visual Preference Task (persons and faces in the stimulus material have been obscured for the preprint version)

### Task 1: Dynamic Visual Preference

The dynamic geometrical images (DGI) versus dynamic social images (DSI) paradigm used in in previous eye-tracking studies on Caucasian children [Moore et al., 2018; Pierce, Conant, Hazin, & Stoner, 2011; Pierce et al., 2015] was adapted for use in Chinese children. Participants were presented with a movie consisting of two rectangular areas of interest which portrayed a single Chinese child or adult dancing on one side and a dynamic geometrical image on the other side (30 different pairs of images were each presented for 2 s and in a continuous sequence). A fixation cross was displayed in the center of the screen for 3 s before the start of the task stimuli. The position of the two types of stimuli on the left or right side was counterbalanced.

### Task 2: Abstract Animate Point Visual Exploration

The biological motion visual preferential task was conducted using point-light displaying animate (walking human or cat) and inanimate (randomly moving point-light) videos [Rutherford & Troje, 2012]. On the left and right side of the screen, animate (human or cat) stimuli *vs*. control inanimate (scrambled versions of the human and cat stimuli) were displayed simultaneously (8 pairs of stimuli with a 10 s presentation duration), and the position of the two types of stimuli on the left and right side was counterbalanced. There was a jittered interval (mean=3 s) between each pair of stimuli where a yellow star was displayed in the center of screen to help keep the attention of the children.

### Task 3: Static Visual Preference

Here we adapted previous tasks [Chevallier et al., 2015; Frazier et al., 2018] and presented pairs of static images, one showing a picture of a child with a happy face playing with a toy while looking outwards towards the subject (i.e. sharing their enjoyment) and the other only including the same toy. In this paradigm 20 pairs of stimuli were each shown for 3 seconds. The position of the two types of stimuli on the left and right side was counterbalanced. There was a jittered interval (mean=3 s) between each pair of stimuli where a yellow star was displayed in the center of screen to help keep the attention of the children.

### Procedure and Eye-Tracking Apparatus

For the experiment children were either seated alone or on a caregivers lap in front of the presentation screen. A Tobii TX300 Binocular Eye Tracker used together with Tobii Studio software (version 3.4.8 Tobii, Stockholm, Sweden) to record visual attention and E-prime linked with the Tobii software used for stimulus presentation. The Tobii system used an I-VT fixation filter. The mean of the right and left eyes was used to calculate fixation. The eye-tracking monitor (TFT-LCD; 23″, 1920×1080) had a refresh rate of 60 Hz. Brightness was 100 % and a five-point calibration procedure was used at the beginning of each experiment. The calibration process was repeated if the initial calibration quality was poor. The total duration of the experimental tasks was around 5 min, with short breaks in between each task.

### Analysis

Social attention was quantified as the mean proportion of total fixation time children spent on looking at the social (vs. non-social) stimuli. This is a widely and uniformly reported measure in eye-tracking studies and meta-analyses [Frazier et al., 2018; Meia Chita-Tegmark, 2016]. Additionally, we measured mean proportion of fixation counts and mean individual fixation durations.

In Task 1, the proportion of total fixation time for children in the ASD and TD groups was investigated in each of the two rectangular areas of interest showing the dynamic social (DSI) and geometric (DGI) images. The proportion of the total fixation time on DSI was used as the dependent variable. In accordance with previous studies [Moore et al., 2018; Pierce et al., 2011, 2015], the dependent variable was calculated as follows: Total looking time = time spent on DSI + time spent on DGI; % total fixation time on social stimuli = (time spent on DSI/Total looking time for DSI and DGI) × 100. Additionally, we measured the proportion of fixation counts and mean fixation durations between the different areas of interest. Children’s data was only included for analysis in this paradigm where they spent >40% of the time looking at the areas of interest [see Frazier et al., 2018].

In Task 2, the areas of interest were the two rectangular regions displaying the animate and inanimate light point videos. The proportion of the total fixation time on human stimuli (HBM) and cat stimuli (CBM) were calculated as follows: % of total fixation time on nonsocial animation = time spent on the cat animation/ (time spent on the cat animation + time spent on the scrambled cat stimulus control) × 100; % of total fixation time on social animation= time spent on the human animation/ (time spent on the human animation + time spent on the scrambled human stimulus control) × 100. Fixation counts and durations were also measured. For this task children needed to spend >15% of the time looking at the areas of interest to be included in the analysis, as in previous studies [Chawarska, Macari & Shic 2013].

In Task 3, the two rectangular area of interest regions included the boy playing with toy region (Social) and toy only (Nonsocial) one. Additionally, we also calculated fixation time on the boy’s face (Face) and the toy in the picture with the boy (Toy1) and the toy in the picture without the boy (Toy2). The proportion of the total fixation times were calculated as follows: % total fixation time on boy playing toy = (time spent on the boy playing with toy / total looking time on both rectangular areas of interest) × 100; % total fixation time on boy’s face = (time spent on face/ time spent at both rectangular areas of interest) × 100; % total fixation time on toy with boy = (time spent on the toy with boy / total looking time on both rectangular areas of interest) × 100; % total fixation to toy without boy = (time spent on the toy without boy/ total looking time on both rectangular areas of interest) × 100. In addition, % time looking at the boy’s face compared to Toy 1 and % time looking at Toy 1 compared to Toy 2 were calculated. In all cases fixation counts and durations were also measured. For this task children also needed to spend >15% of the time looking at the areas of interest to be included in the analysis, as in previous studies [Chawarska, Macari & Shic 2013].

### Statistical analyses

To compare proportion of total fixation time and fixation counts as well as mean individual fixation durations for DSI, HBM, CBM, Social, Face, Toy1, and Toy2 between the ASD and TD groups, independent T-tests were performed. For Tasks 2 and 3, 2-way ANOVAs were used to compare multiple variables followed by Bonferonni corrected post-hoc comparisons. The association between proportion of total fixation duration and fixation counts and mean individual fixation durations to the different stimuli and ADOS-2 scores was conducted using Pearson correlation analysis. All statistical analyses were performed using SPSS-22. Effect sizes (partial eta-squared, η^2^ p for F statistics and Cohen’s *d* for t-tests) were reported with P-values for significant main effects and interactions. Receiver operating characteristic (ROC) analyses were used for investigating the ability of specific parameters in the three tasks for discriminating between the ASD and TD groups in order to compare the diagnostic utility of each.

## Results

A total of 3 children in the ASD were excluded from analysis due to problems with eye-tracking data collection and in Tasks 2 and 3, 6 and 12 children respectively in the ASD group had to be excluded from analysis due to viewing the screen for < 15% of the time. Following these exclusions ASD children still spent significantly less total time than TD children looking at the display screen (Task 1 total task time = 63s: ASD mean ± SD 56.17 ± 9.12s, TD 61.56 ± 4.22s, t_64_ = 3.05, *p* = 0.004; Task 2 total task time = 101s: ASD 50.81 ± 17.14s, TD 75.81 ± 12.51s, t_61_ = 6.68, *p* < 0.001; Task 3 total task time = 120s: ASD 57.14 ± 14.52s, TD 83.74 ±15.52s, t_55_ = 6.514, *p* < 0.001). Overall, children in both groups spent proportionately more time looking at the display screen in Task 1 than in Tasks 2 and 3. In Tasks 2 and 3 during the fixation periods between stimuli children in both groups were particularly less attentive to the screen.

### Visual Preference Pattern Differences between ASD and TD groups and associations with symptom severity

#### Task 1

Figure 2 shows that there was a significant difference in the proportion of total fixation time on the dynamic social images (DSI) between the two groups (ASD *vs*. TD, t_64_ = 3.85, *p*<0.001 Cohen’s d=0.96). Whereas TD children spent equivalent proportions of time looking at DSI and DGI (53.4% on DSI and 47.6% on DGI) those with ASD spent a greater proportion of time looking at DGI compared with DSI (35.3% on DSI and 64.7% on DGI) (see Fig.2A). There was also a significant difference between ASD and TD groups for the proportion of fixation counts to the DSI as opposed to DGI (t_64_ = 4.24, *p* < 0.001 Cohen’s *d* = 1.07) due to the ASD group showing proportionately less fixations than the TD group for DSI (ASD: 41.9% on DSI and 58.1% on DGI; TD 59.2% on DSI and 41.8% on DGI) (see Fig. 2C). There was no significant difference between the groups for mean individual fixation durations on DSI (*p* >0.130).

**Fig.2.**
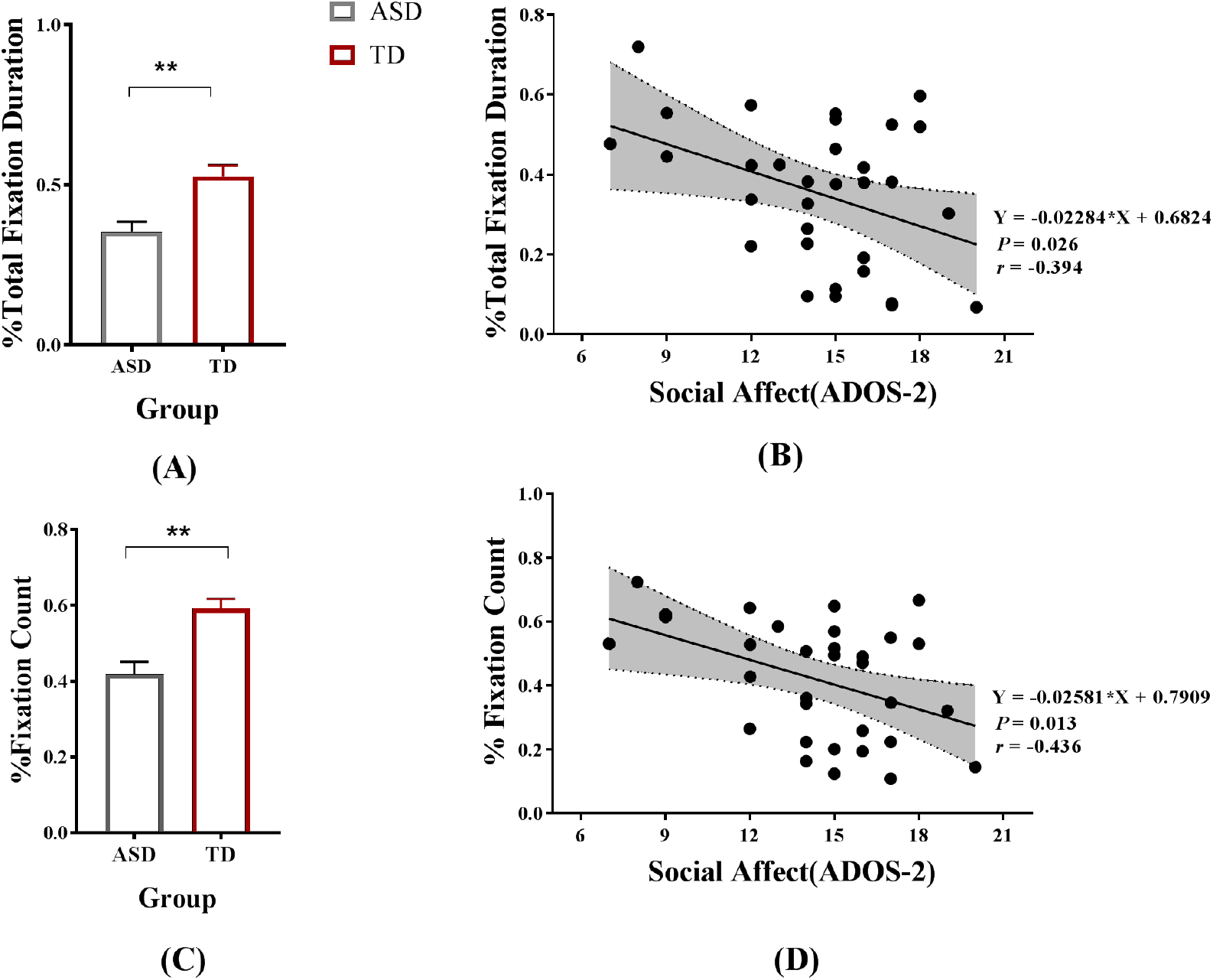
Differences between ASD and TD children in Task 1 (A) Group difference of the proportion of total fixation time on the dynamic social images (DSI). (B) Negative correlation between social affect scores (ADOS-2) and total fixation time. (C) and (D) the same information for proportion of fixation counts, p<0.05*, p<0.01**, two-tailed t-test. Bars indicate M ± SE.

There was a significant association between the proportion of total fixation time on the DSI and ADOS-2 social affect scores (SA) (*r* = −0.40, *p* = 0.026) (see Fig.2B) but not with the total ADOS-2 score (*r* =-0.20, *p* = 0.270) or Restricted and Repetitive Behavior score (RRB) (*r* =0.17, *p* = 0.339). There was also a significant negative correlation between the proportion of fixation counts for DSI in children in the ASD group and their ADOS-2 SA scores (*p* = 0.013, r = −0.44) (see Fig.2D) but not with their total ADOS-2 (*r* =-0.27, *p* = 0.133) or RRB scores (*r* =0.10, *p* = 0.568).

#### Task 2

For proportion of total fixation time a two-way ANOVA with group and stimulus type (cat vs. human) revealed significant main effects of stimulus type (F_1, 61_ = 4.799, *p* = 0.032, η^2^ *p* = 0.073) and group (F_1, 61_ = 5.333, *p* = 0.024, η^2^ *p* = 0.080) but no significant interaction (F_1, 61_ = 2.436, *p* = 0.124, η^2^ *p* = 0.038). An exploratory post-hoc analysis showed a significant difference for the proportion of time looking at the cat animation between ASD and TD groups (*p*= 0.010, Cohen’s *d* =0.68) with the ASD group showing a greater visual preference (M ± SD, 52.18±15.63%) than the TD group (M ± SD, 42.99±11.90%). While there was a significant difference between the proportion of time looking at the cat as opposed to the human animation in the TD group (*p*= 0.008, Cohen’s *d* =0.67) this was not the case in the ASD group (*p*= 0.670), with the TD group showing a greater preference for viewing the human animation (M ± SD, human: 52.18±15.63%; cat: 42.99±11.90%) (see Fig. 3). For proportion of fixation counts a two-way ANOVA revealed significant main effect of group (F_1, 61_ =5.966, *p* = 0.017, η^2^ *p* = 0.089), but not for stimulus type (F_1, 61_ = 0.00, *p* = 0.994) and no significant interaction (F_1, 61_ = 0.172, *p* = 0.680). An exploratory post-hoc analysis showed a marginal difference for the proportion of fixation counts at the cat (*p* = 0.052) and human (*p* = 0.093) animations between ASD and TD groups, with the ASD group tending to have a higher fixation count than the TD group.

**Fig.3.**
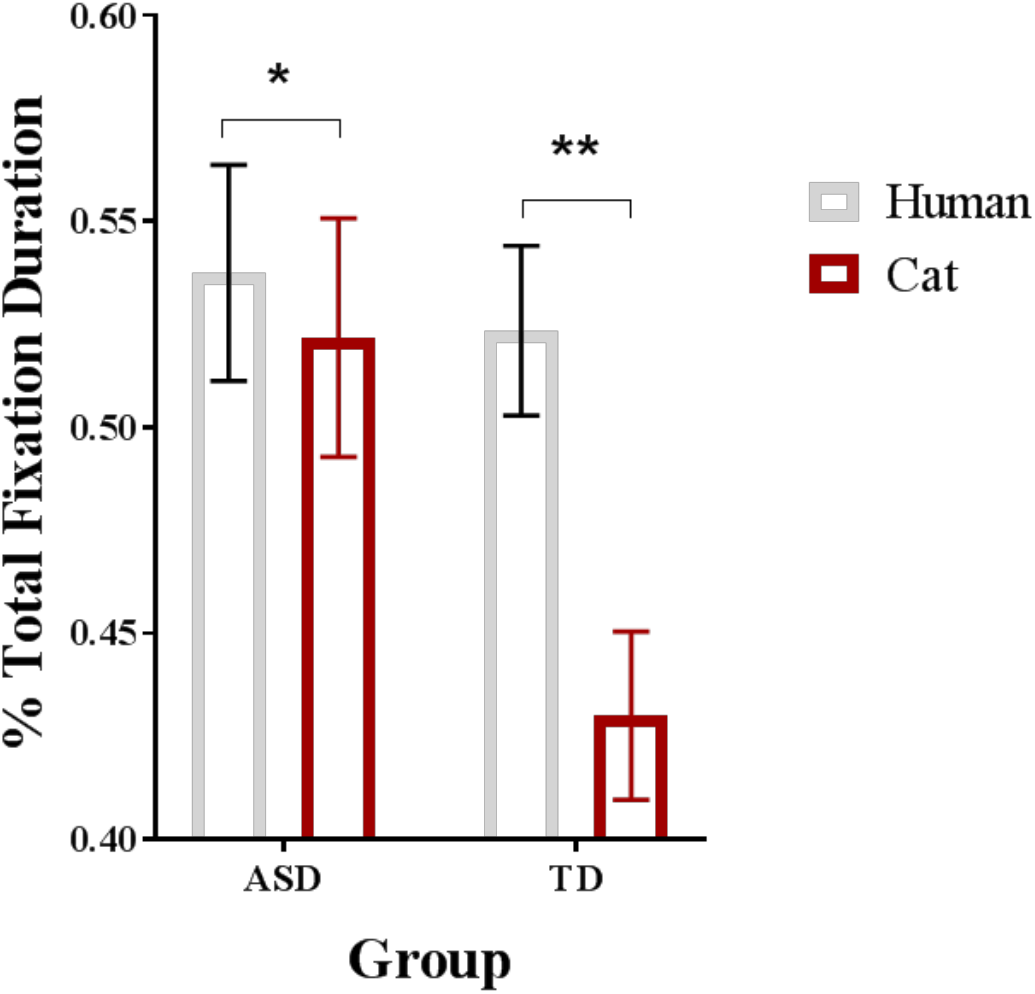
Differences between ASD and TD children in Task 2. Group differences for the proportion of total fixation time on animated human and cat stimuli compared with scrambled controls. p<0.05*, p<0.01**, two-tailed t-test. Bars indicate M ± SE.

There was no significant group difference of mean individual fixation durations for cat or human animations and their scrambled controls (*ps*>0.114).

There was no significant correlation found between and of the eye tracking measures and symptom severity (SA, RRB and ADOS-2 total scores) in the ASD group (all *rs* <0.234, *ps* >0.223).

#### Task 3

For the proportion of fixation time a two-way ANOVA with group and stimuli (4 different AOIs: Social, Face, Toy1, Toy2) as factors revealed significant main effects of stimuli (F_1, 55_ = 103.918, *p* < 0.001, η^2^ *p* = 0.5654) and group (F_1, 55_ = 15.615, *p* < 0.001, η^2^ *p* = 0.221) but no significant interaction (F_1, 55_ = 0.630, *p* =0.597, η^2^ *p* = 0.011). An exploratory post-hoc analysis showed a significant group difference for time looking at Toy1 (*p*= 0.018, Cohen’s *d* =0.67) with the TD group showing a greater proportion (M ± SD, 27.26±7.41%) than the ASD one (M ± SD, 22.23±8.03%). There were no significant group differences for the proportion of time spent looking at the social picture (*p* = 0.105), the face alone in the social picture (*p* = 0.105) or Toy2 (*p* = 0.291) (see Fig. 4). Additional group comparisons between the proportion of time spent looking at Face *vs* Toy1 and between Toy1 *vs*. Toy 2 were also made but revealed no significant differences (all *ps* > 0.277).

**Fig.4.**
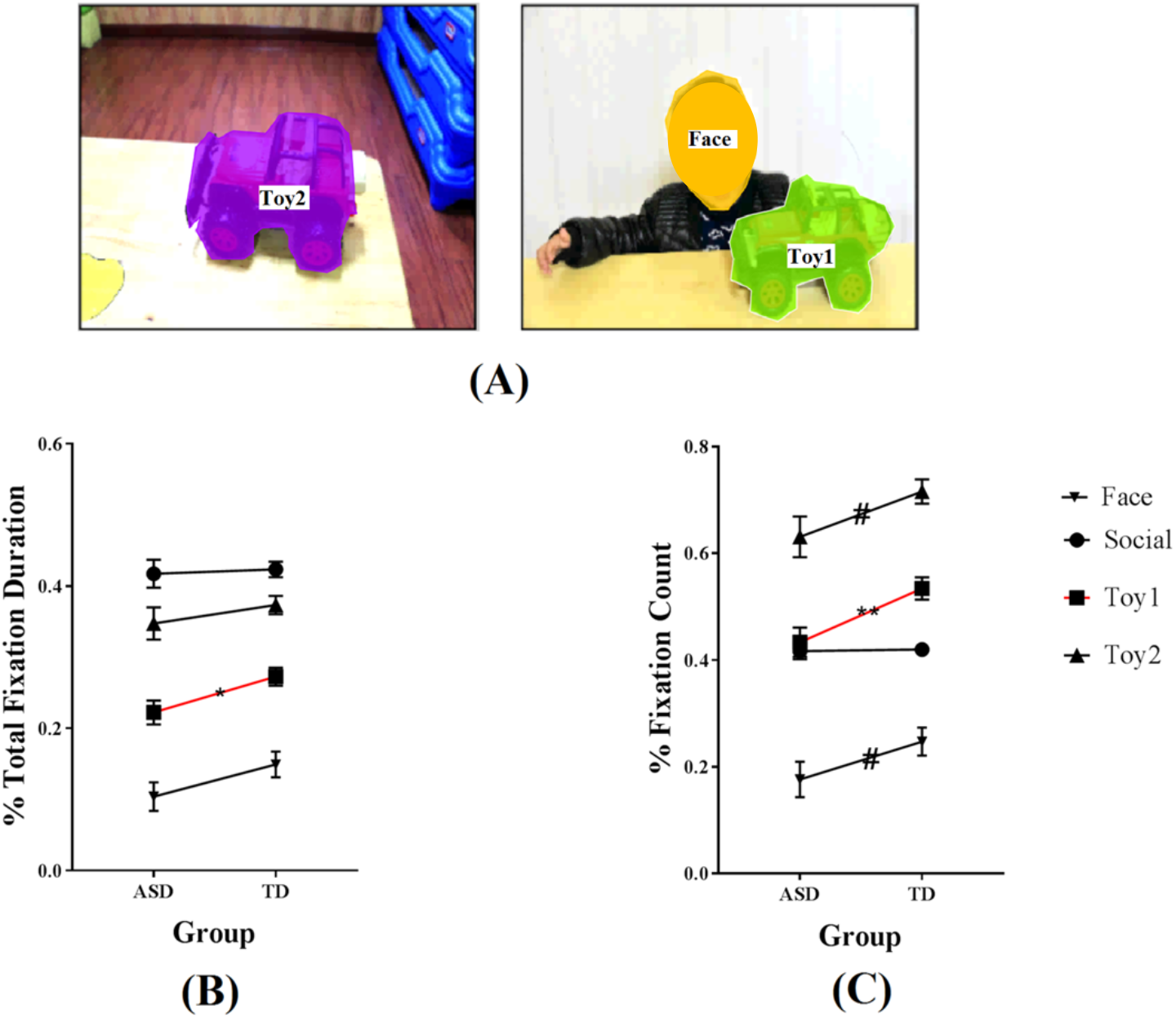
Differences between ASD and TD children in Task 3 (A) Pictures show regions of interest used in Task 3 (B) Group differences in the proportion of total fixation time on the four different regions of interest in Task 3. (C) Group differences in the proportion of fixation counts on the four different regions of interest in Task 3.

For the proportion of fixation counts a two-way ANOVA revealed significant main effects of stimuli (F_1, 55_ =98.850, *p* < 0.001, η^2^ *p* = 0.643) and group (F_1, 55_ = 30.417, *p* < 0.001, η^2^ *p* = 0.356) but no significant interaction (F_1, 55_ = 1.258, *p*=0.291, η^2^ *p* = 0.022). An exploratory post-hoc analysis revealed no significant group difference for Social, Face and Toy2 (*ps*>0.06) but a significant group difference for Toy1 (*p*= 0.004, Cohen’s *d* =2.04), with the TD group showing a greater proportion (M ± SD, 41.24±11.14) than the ASD group (M ± SD, 21.26±7.91). Additional group comparisons between the proportion of fixation counts for Face *vs* Toy1 and between Toy1 *vs*. Toy 2 were also made but revealed no differences (all *ps* > 0.29).

There was no significant group difference in mean individual fixation durations for any of the areas of interest (Social, Face, Toy1, Toy2) (*ps*>0.126).

The correlations between eye-tracking measures of visual preference for Toy1 and symptom severity in the ASD group were not significant (SA, RRB and ADOS total score) (all *rs* <0.264, *ps* >0.223).

### Relative diagnostic utility of the three task paradigms

The diagnostic utility for distinguishing ASD from TD children for the three different tasks was assessed by performing a receiver operating characteristic (ROC) analysis. For Task 1 the proportion of total fixation time for DSI revealed the area under the ROC curve was 0.743 (95% CI 0.625-0.860, *p* = 0.001). For Task 2 the proportion of total fixation time for the cat animation revealed the area under the ROC curve was 0.691 (95% CI 0.553-0.829, *p* = 0.010) and for Task 3 the proportion of total fixation time for Toy1 revealed an area under the ROC curve of 0.687 (95% CI 0.543-0.831, *p* = 0.018) (see Fig.5). For Task 1 the proportion of fixation counts for DSI revealed an area under the ROC curve of 0.764 (95% CI 0.651-0.877, *p*<0.001). For Task 2 the proportion of fixation duration for the cat animation revealed an area under the ROC curve of 0.640 (95% CI 0.497-0.783, *p* = 0.057) and for Task 3 the proportion of total fixation time for Toy1 revealed an area under the ROC curve of 0.710 (95% CI 0.569-0.851, *p* = 0.008) (see Fig. 5).

**Fig.5.**
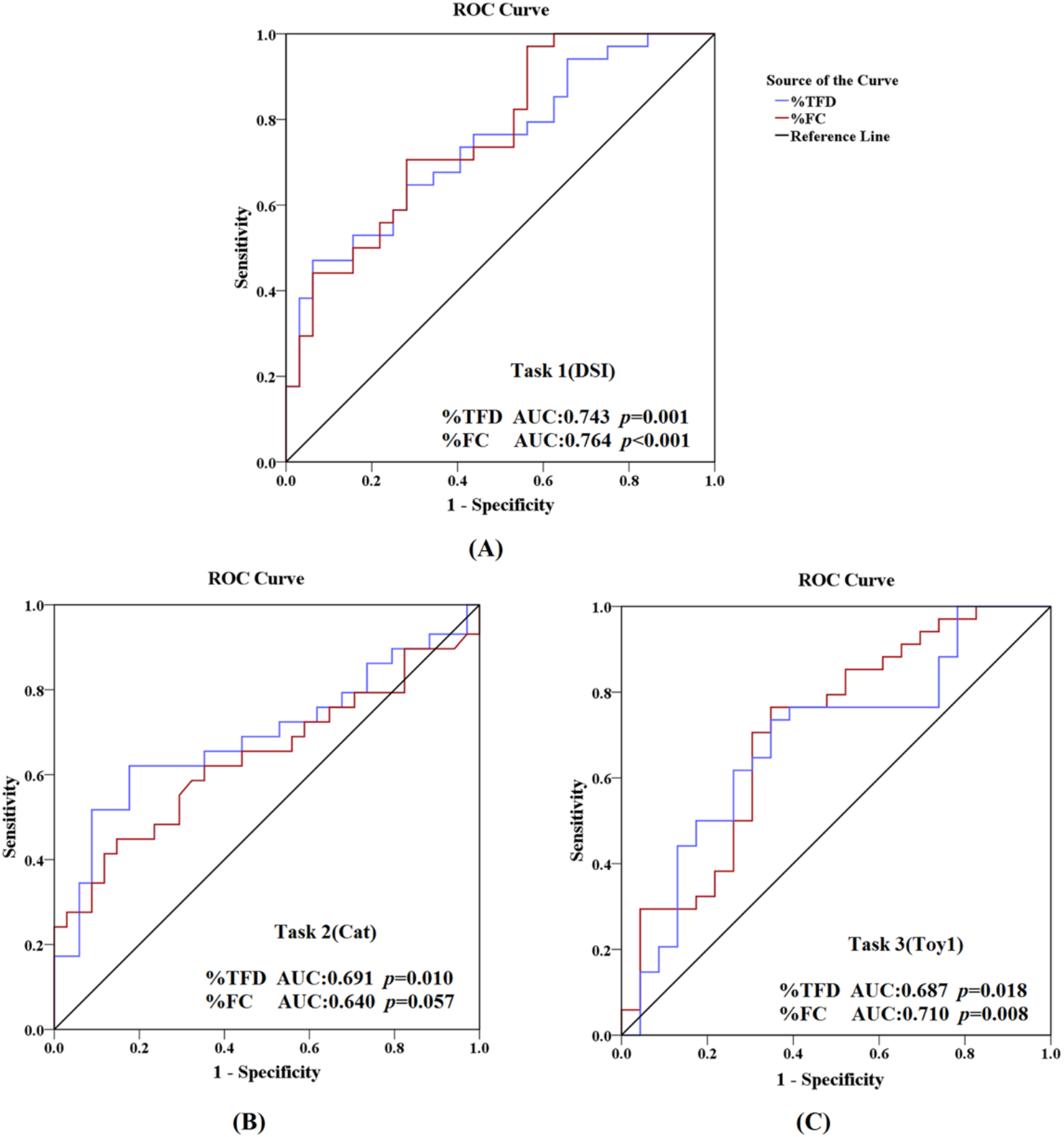
Receiver operating curves (ROC) for discrimination of ASD from TD children for proportion of total fixation time (%TFD) and fixation count (%FC) for (A) DSI in Task1 (B) Cat animation in Task 2 and (C) Toy 1 in Task 3.

## Discussion

The main objective of the current study was to compare and contrast the relative efficacies of three different, commonly used eye-tracking paradigms firstly for discriminating between Chinese ASD and TD children and secondly to show a significant association with symptom severity as measured using ADOS-2. Overall our results show that a Chinese version of the dynamic dancing individuals versus dynamic geometric patterns originally developed by Karen Pierce and colleagues (Pierce et al., 2011; 2015) was the most effective both for showing differences between ASD and TD children (with both large effect sizes and areas under the ROC curve) and for demonstrating associations with the severity of social affect symptoms. Importantly as well, all the ASD children paid attention to the display screen for almost the whole of the period of the task whereas in the other two tasks used many of them did not and had to be excluded. As such this specific dynamic task may be particularly suitable and sensitive for aiding both diagnosis and assessing social affect symptom severity across cultures.

In line with many previous studies [Moore et al., 2018; Pierce et al., 2011, 2015] children in the ASD group generally spent less time looking at the display screen than ones in the TD group and thus the most effective measure for group comparisons is the relative proportion of time spent viewing one stimulus compared with another. Indeed, in accordance with the original definition autistic children tend to shut out anything that comes to them from outside [Kanner, 1943], and therefore pay less, or an altered pattern of, visual attention to environment cues (including social and nonsocial) than TD children. Autistic individuals also tend to exhibit an enhanced self-focus [Burrows, Laird, & Uddin, 2016; Vasudeva & Hollander, 2017], it can result in reduced external attention. In the dynamic dancing versus geometric image task none of the ASD children needed to be excluded for insufficient attention (and with a higher threshold of >40%) to the display screen with the remainder looking at the screen on average for 89% of the time compared with 98% in TD children. After exclusions in the other two tasks (and with a lower threshold of >15%) ASD children only spent 50.3% (Task 2) and 45.6% (Task 3) of the time looking at the screen compared with 75.1% and 69.8% in TD children. In these latter two tasks inattention was particularly during the jittered fixation intervals between stimuli which are not included in Task 1, and this suggests that such routine fixation periods may need to be minimized when using eye-tracking protocols in ASD children. However, the dynamic dancing versus geometric stimuli task is clearly very effective in gaining the attention of both ASD and TD children, which further underlines its utility.

In agreement with previous studies on Caucasian children (Franchini et al., 2017; Moore et al., 2018; Pierce et al., 2011; 2015) in the Dynamic Visual Preference Task ASD children show a greater attentional preference for dynamic geometrical patterns compared with individuals dancing/exercising whereas in TD children the opposite is the case. The strong association we found between the reduced interest in the dynamic social stimuli and the severity of social deficits in ASD is also in agreement with studies in Caucasian children. Importantly, both of these findings are not restricted to specific stimulus sets since those we used were different to ones in other previous studies. In the current study both the proportion of total fixation duration and also of fixation counts were sensitive measures but not mean durations of individual fixations. Thus children with ASD spend proportionately less time looking at the DSI because they show a reduced number of fixations to them rather and a reduced duration of individual fixations. It is possible that specific types of dynamic geometrical images may be more attractive than others to ASD individuals and that varying the number of individuals and type of movements they exhibit might also increase their attractiveness to TD individuals. A small-scale study for example found that displaying multiple dancing individuals was more effective than only one [Shi et al., 2015]. If so, then additional refinements to this paradigm might serve to further increase its diagnostic sensitivity and utility.

While a number of previous studies have reported than ASD individuals show reduced attention towards biological motion compared with scrambled motion [Annaz et al., 2012; Falck-Ytter et al., 2013; Wang et al., 2015] we did not replicate this in the present study. Indeed, ASD children were generally attracted by biological motion and even significantly more so that TD ones for the cat stimulus. Although this latter group different had a reasonable ability to discriminate between ASD and TD children (ROC area under the curve of 0.69) it was not associated with symptom severity. The only other difference was that TD children were significantly more interested in human than cat stimuli whereas ASD ones showed equivalent interest in them. Our findings with this paradigm are in line with a recent study that individuals with ASD can be more sensitive to the biological motion and both ASD and the TD groups are able to detect the perceptual information [Rosanna et al., 2018]. Many other studies have also failed to demonstrate significant differences between ASD and TD children using versions of this biological motion paradigm [Atkinson, 2009; Jones et al., 2011; Koldewyn, Whitney & Rivera, 2010; Murphy et al., 2009; Saygin, Cook, & Blakemore, 2010]. Thus, our overall conclusion is that this protocol in its current format may not be sufficiently robust for use as an ASD marker.

Our findings using an adapted version of Static Visual Preference Task (Task 3) indicated that although there was no visual preference difference between the two groups for the face of the child playing with the toy, the ASD group spent proportionately less time looking at the toy the boy was holding in his hand. This could indicate that ASD children were less motivated to share the interest of the child in the picture for the toy. This pattern of altered preference also showed a reasonable ability to discriminate between ASD and TD individuals (ROC area under the curve of 0.67), however as with the biological motion paradigm findings it showed no significant association with symptom severity. Furthermore, many ASD children failed to pay sufficient attention to the screen during the paradigm which make it difficult to use as a general diagnostic aid.

In summary, our findings comparing the efficacy of three different eye-tracking paradigms as a marker for ASD in young Chinese children, together with the severity of its social symptoms have emphasized the utility of the dynamic paradigm comparing a dancing human with geometric patterns. Importantly, this paradigm appears to be effective both across cultures and stimulus sets and possibly further research may be able to further refine its utility through choice of specific stimulus content and potentially also for use as a diagnostic aid in younger children at risk of ASD.

## References

American Psychiatric Association. (1994). Diagnostic and statistical manual of mental disorders (4th ed.). Washington DC: American Psychiatric Association Publishing.

Annaz, D., Campbell, R., Coleman, M., Milne, E., & Swettenham, J. (2012). Young children with autism spectrum disorder do not preferentially attend to biological motion. Journal of Autism and Developmental Disorders, 42(3), 401–408.

Atkinson, A. P. (2009). Impaired recognition of emotions from body movements is associated with elevated motion coherence thresholds in autism spectrum disorders. Neuropsychologia, 47(13), 3023–3029.

Brannan, A. M., Heflinger, C. A., & Bickman, L. (1997). The Caregiver Strain Questionnaire: Measuring the impact on the family of living with a child with serious emotional disturbance. Journal of Emotional and Behavioral Disorders, 5(4), 212–222.

Burrows, C. A., Laird, A. R., & Uddin, L. Q. (2016). Functional connectivity of brain regions for self- and other-evaluation in children, adolescents and adults with autism. Developmental Science, 19(4), 564–580.

Chevallier, C., Parish-Morris, J., Mcvey, A., Rump, K. M., Sasson, N. J., Herrington, J. D., & Schultz, R. T. (2015). Measuring social attention and motivation in autism spectrum disorder using eye-tracking: Stimulus type matters. Autism Research, 8(5), 620–628.

Chita-Tegmark, M. (2016). Social attention in ASD: a review and meta-analysis of eye-tracking studies. Research in developmental disabilities, 48, 79–93.

Rubenstein, E., Schieve, L., Wiggins, L., Rice, C., Braun, K. V. N., Christensen, D., Durkind M., Daniels J., & Lee, L. C. (2018). Trends in documented co-occurring conditions in children with autism spectrum disorder, 2002–2010. Research in developmental disabilities, 83, 168–178.

Constantino, J. N., & Gruber, C. P. (2012). The Social Responsiveness Scale (2nd.) (SRS-2) [Manual]. Torrance, CA: Western Psychological Services.

Donald, A. K. W. J. R. S. F. V. (2002). Visual Fixation Patterns During Viewing of Naturalistic Social Situations as Predictors of Social Competence in Individuals With Autism. Matters of Conflict: Material Culture, Memory and the First World War, 59, 149–165.

Edey, R., Cook, J., Brewer, R., Bird, G., & Press, C. (2018). Adults with autism spectrum disorder are sensitive to the kinematic features defining natural human motion. Autism Research. doi: 10.1002/aur.2052

Falck-Ytter, T., Bölte, S., & Gredebäck, G. (2013). Eye tracking in early autism research. Journal of Neurodevelopmental Disorders, 5(1), 28.

Franchini, M., Glaser, B., De Wilde, H. W., Gentaz, E., Eliez, S., & Schaer, M. (2017). Social orienting and joint attention in preschoolers with autism spectrum disorders. PLoS ONE, 12(6).

Frazier, T. W., Klingemier, E. W., Parikh, S., Speer, L., Strauss, M. S., Eng, C., Hardan A.Y., & Youngstrom, E. A. (2018). Development and Validation of Objective and Quantitative Eye Tracking-Based Measures of Autism Risk and Symptom Levels. Journal of the American Academy of Child & Adolescent Psychiatry, 57(11), 858–866.

Fujisawa, T. X., Tanaka, S., Saito, D. N., Kosaka, H., & Tomoda, A. (2014). Visual attention for social information and salivary oxytocin levels in preschool children with autism spectrum disorders: an eye-tracking study. Frontiers in neuroscience, 8, 295.

Jones, C. R. G., Swettenham, J., Charman, T., Marsden, A. J. S., Tregay, J., Baird, G., Simonoff, E., & Happé, F. (2011). No evidence for a fundamental visual motion processing deficit in adolescents with autism spectrum disorders. Autism Research, 4(5), 347–357.

Kanne, S. M., Randolph, J. K., & Farmer, J. E. (2008). Diagnostic and assessment findings: A bridge to academic planning for children with autism spectrum disorders. Neuropsychology Review, 18(4), 367–384.

Kanner, L. (1943). Autistic disturbances of affective contact. Nervous child, 2(3), 217–250.

Klin, A., Lin, D. J., Gorrindo, P., Ramsay, G., & Jones, W. (2009). Two-year-olds with autism orient to non-social contingencies rather than biological motion. Nature, 459(7244), 257–261.

Kim, Y. S., Leventhal, B. L., Koh, Y. J., Fombonne, E., Laska, E., Lim, E. C., & Song, D. H. (2011). Prevalence of autism spectrum disorders in a total population sample. American Journal of Psychiatry, 168(9), 904–912.

Koldewyn, K., Whitney, D., & Rivera, S. M. (2010). The psychophysics of visual motion and global form processing in autism. Brain, 133(2), 599–610.

Lam, K. S., & Aman, M. G. (2007). The Repetitive Behavior Scale-Revised: independent validation in individuals with autism spectrum disorders. Journal of autism and developmental disorders, 37(5), 855–866.

Lord, C., Elsabbagh, M., Baird, G., & Veenstra-Vanderweele, J. (2018). Autism spectrum disorder. The Lancet, 0(0), 508–520.

Lord, C., Luyster, R. J., Gotham, K., Guthrie, W. (2012). Autism diagnostic observation schedule, second edition (ADOS-2) manual (Part II): Toddler module. Torrance, CA: Western Psychological Services.

Lord, C., Rutter, M., DiLavore, P. C., Risi, S., Gotham, K., & Bishop, S. L. (2012). Autism Diagnostic Observation Schedule, second edition (ADOS-2) [Manual]. Torrance, CA: Western Psychological Services.

Moore, A., Wozniak, M., Yousef, A., Barnes, C. C., Cha, D., Courchesne, E., & Pierce, K. (2018). The geometric preference subtype in ASD: Identifying a consistent, early-emerging phenomenon through eye tracking. Molecular Autism, 9(1), 1–13.

Murphy, P., Brady, N., Fitzgerald, M., & Troje, N. F. (2009). No evidence for impaired perception of biological motion in adults with autistic spectrum disorders. Neuropsychologia, 47(14), 3225–3235.

Pierce, K., Marinero, S., Hazin, R., Mckenna, B., & Barnes, C. C. (2015). Archival Report Eye Tracking Reveals Abnormal Visual Preference for Geometric Images as an Early Biomarker of an Autism Spectrum Disorder Subtype Associated with Increased Symptom Severity. Biological Psychiatry, (9), 1–10.

Rutter, M., Bailey, A., & Lord, C. (2003). The social communication questionnaire: Manual. Western Psychological Services.

Rutherford, M. D., & Troje, N. F. (2012). IQ predicts biological motion perception in autism spectrum disorders. Journal of Autism and Developmental Disorders, 42(4), 557–565.

Rutter, M., Le Couteur, A., & Lord, C. (2003). Autism diagnostic interview-revised. Los Angeles, CA: Western Psychological Services, 29, 30.

Saygin, A. P., Cook, J., & Blakemore, S. J. (2010). Unaffected perceptual thresholds for biological and non-biological form-from-motion perception in autism spectrum conditions. PLoS ONE, 5(10), 1–7.

Shi, L., Zhou, Y., Ou, J., Gong, J., Wang, S., Cui, X., Lyu, H., Zhao, J., & Luo, X. (2015). Different visual preference patterns in response to simple and complex dynamic social stimuli in preschool-aged children with autism spectrum disorders. PLoS ONE, 10(3), 1–16.

Tick, B., Bolton, P., Happé, F., Rutter, M., & Rijsdijk, F. (2016). Heritability of autism spectrum disorders: A meta-analysis of twin studies. Journal of Child Psychology and Psychiatry and Allied Disciplines, 57(5), 585–595.

Vasudeva, S. B., & Hollander, E. (2017). Body dysmorphic disorder in patients with autism spectrum disorder: A reflection of increased local processing and self-focus. American Journal of Psychiatry, 174(4), 313–316.

Wang, L., Chien, S. H., Hu, S., Chen, T., & Chen, H. (2015). Research in Autism Spectrum Disorders Children with autism spectrum disorders are less proficient in action identification and lacking a preference for upright point-light biological motion displays. Research in Autism Spectrum Disorders, 11(91), 63–76.

Wang, F., Lu, L., Wang, S., Zhang, L., Ng, C. H., Ungvari, G. S., Cao, X., Lu, J., Hou, C., Jia, F., & Xiang, Y. (2018). The prevalence of autism spectrum disorders in China: a comprehensive meta-analysis. International journal of biological sciences, 14(7), 717.

World Health Organization. (1994). International classification of diseases (10th ed.). Geneva, Switzerland: Author.

